# Reagent contamination can critically impact sequence-based microbiome analyses

**DOI:** 10.1101/007187

**Authors:** Susannah J Salter, Michael J Cox, Elena M Turek, Szymon T Calus, William O Cookson, Miriam F Moffatt, Paul Turner, Julian Parkhill, Nick Loman, Alan W Walker

## Abstract

The study of microbial communities has been revolutionised in recent years by the widespread adoption of culture independent analytical techniques such as 16S rRNA gene sequencing and metagenomics. One potential confounder of these sequence-based approaches is the presence of contamination in DNA extraction kits and other laboratory reagents. In this study we demonstrate that contaminating DNA is ubiquitous in commonly used DNA extraction kits, varies greatly in composition between different kits and kit batches, and that this contamination critically impacts results obtained from samples containing a low microbial biomass. Contamination impacts both PCR based 16S rRNA gene surveys and shotgun metagenomics. These results suggest that caution should be advised when applying sequence-based techniques to the study of microbiota present in low biomass environments. We provide an extensive list of potential contaminating genera, and guidelines on how to mitigate the effects of contamination. Concurrent sequencing of negative control samples is strongly advised.

## INTRODUCTION

Culture-independent studies of microbial communities are revolutionising our understanding of microbiology and revealing exquisite interactions between microbes, animals and plants. Two widely used techniques are deep sequence surveying of PCR-amplified marker genes such as 16S rRNA, or whole-genome shotgun metagenomics, where the entire complement of community DNA is sequenced *en masse*. While both of these approaches are powerful, they have important technical caveats and limitations, which may distort the taxonomic distribution and frequencies observed in the sequence dataset. Such limitations, which have been well reported in the literature, include choices relating to sample collection, sample storage and preservation, DNA extraction, amplifying primers, sequencing technology, read length and depth and bioinformatics analysis techniques^1,2^.

A related additional problem is the introduction of contaminating microbial DNA during sample preparation. Possible sources of DNA contamination include molecular biology grade water^3–9^, PCR reagents^10–15^ and DNA extraction kits themselves^16^. Contaminating sequences matching water- and soil-associated bacterial genera including *Acinetobacter, Alcaligenes, Bacillus, Bradyrhizobium, Herbaspirillum, Legionella, Leifsonia, Mesorhizobium, Methylobacterium, Microbacterium, Novosphingobium, Pseudomonas, Ralstonia, Sphingomonas, Stenotrophomonas* and *Xanthomonas* have been reported previously^3,15,17,18^. The presence of contaminating DNA is a particular challenge for researchers working with samples containing a low microbial biomass. In these cases, the low amount of starting material may be effectively swamped by the contaminating DNA and generate misleading results.

Although the presence of such contaminating DNA has been reported in the literature, usually associated with PCR-based studies, its possible impact on high-throughput 16S rRNA gene-based profiling and shotgun metagenomics studies has not been reported. In our laboratories we routinely sequence negative controls, consisting of “blank” DNA extractions and subsequent PCR amplifications. Despite adding no sample template at the DNA extraction step, these negative control samples often yield a range of contaminating bacterial species (see Table 1), which are often also visible in the human-derived samples that are processed concomitantly with the same batch of DNA extraction kits. The presence of contaminating sequences is greater in low-biomass samples (such as from the blood or lung) than in high-biomass samples (such as from faeces), suggesting that there is a critical tipping point where contaminating DNA becomes dominant in sequence libraries.

**Table 1.**
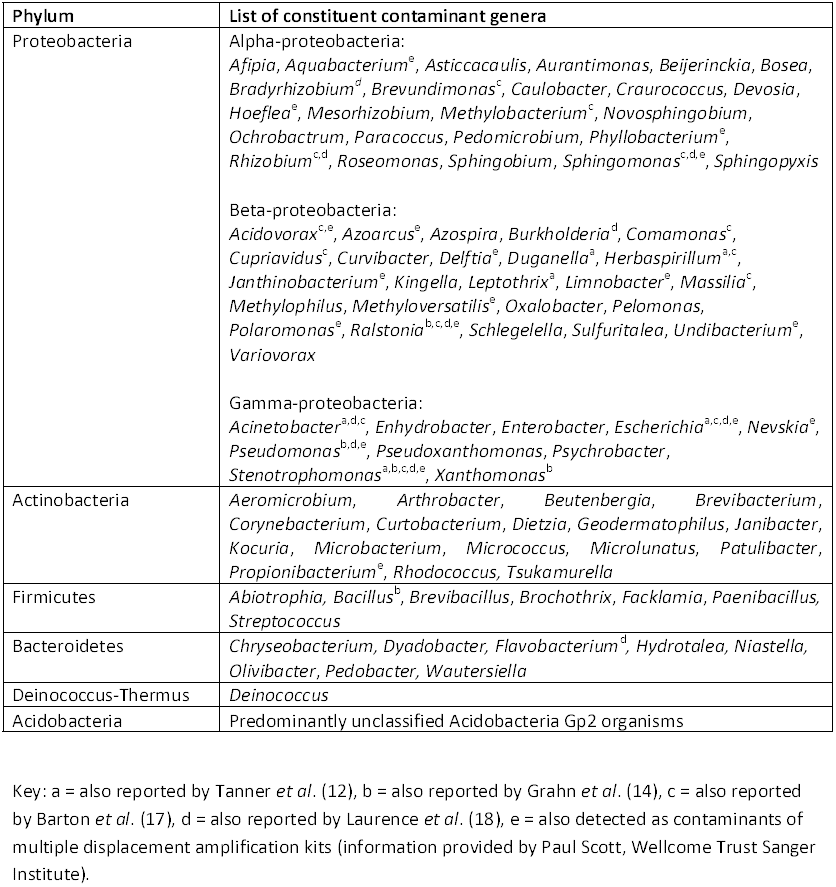
List of contaminant genera detected in sequenced negative “blank” controls

| Phylum | List of constituent contaminant genera |
| --- | --- |
| Proteobacteria | Alpha-proteobacteria:<br>*Afipia*, *Aquabacterium*<sup>e</sup>, *Asticcacaulis*, *Aurantimonas*, *Beijerinckia*, *Bosea*, *Bradyrhizobium*<sup>d</sup>, *Brevundimonas*<sup>c</sup>, *Caulobacter*, *Craurococcus*, *Devosia*, *Hoeflea*<sup>e</sup>, *Mesorhizobium*, *Methylobacterium*<sup>c</sup>, *Novosphingobium*, *Ochrobactrum*, *Paracoccus*, *Pedomicrobium*, *Phyllobacterium*<sup>e</sup>, *Rhizobium*<sup>c,d</sup>, *Roseomonas*, *Sphingobium*, *Sphingomonas*<sup>c,d,e</sup>, *Sphingopyxis*<br>Beta-proteobacteria:<br>*Acidovorax*<sup>c,e</sup>, *Azoarcus*<sup>e</sup>, *Azospira*, *Burkholderia*<sup>d</sup>, *Comamonas*<sup>c</sup>, *Cupriavidus*<sup>c</sup>, *Curvibacter*, *Delftia*<sup>e</sup>, *Duganella*<sup>a</sup>, *Herbaspirillum*<sup>a,c</sup>, *Janthinobacterium*<sup>e</sup>, *Kingella*, *Leptothrix*<sup>a</sup>, *Limnobacter*<sup>e</sup>, *Massilia*<sup>c</sup>, *Methylophilus*, *Methyloversatilis*<sup>e</sup>, *Oxalobacter*, *Pelomonas*, *Polaromonas*<sup>e</sup>, *Ralstonia*<sup>b,c,d,e</sup>, *Schlegelella*, *Sulfuritalea*, *Undibacterium*<sup>e</sup>, *Variovorax*<br>Gamma-proteobacteria:<br>*Acinetobacter*<sup>a,d,c</sup>, *Enhydrobacter*, *Enterobacter*, *Escherichia*<sup>a,c,d,e</sup>, *Nevskia*<sup>e</sup>, *Pseudomonas*<sup>b,d,e</sup>, *Pseudoxanthomonas*, *Psychrobacter*, *Stenotrophomonas*<sup>a,b,c,d,e</sup>, *Xanthomonas*<sup>b</sup> |
| Actinobacteria | *Aeromicrobium*, *Arthrobacter*, *Beutenbergia*, *Brevibacterium*, *Corynebacterium*, *Curtobacterium*, *Dietzia*, *Geodermatophilus*, *Janibacter*, *Kocuria*, *Microbacterium*, *Micrococcus*, *Microlunatus*, *Patulibacter*, *Propionibacterium*<sup>e</sup>, *Rhodococcus*, *Tsukamurella* |
| Firmicutes | *Abiotrophia*, *Bacillus*<sup>b</sup>, *Brevibacillus*, *Brochothrix*, *Facklamia*, *Paenibacillus*, *Streptococcus* |
| Bacteroidetes | *Chryseobacterium*, *Dyadobacter*, *Flavobacterium*<sup>d</sup>, *Hydrotalea*, *Niastella*, *Olivibacter*, *Pedobacter*, *Wautersiella* |
| Deinococcus-Thermus | *Deinococcus* |
| Acidobacteria | Predominantly unclassified Acidobacteria Gp2 organisms |

Key: a = also reported by Tanner *et al.* (12), b = also reported by Grahn *et al.* (14), c = also reported by Barton *et al.* (17), d = also reported by Laurence *et al.* (18), e = also detected as contaminants of multiple displacement amplification kits (information provided by Paul Scott, Wellcome Trust Sanger Institute).

Many recent publications^19–37^ describe important or core microbiota members, often members that are biologically unexpected, and which overlap with previously-described contaminant genera. Spurred by this, and the results from negative control samples in our own laboratories when dealing with low-input DNA samples, we investigated the impact of contamination on microbiota studies and explored methods to limit the impact of such contamination. In this study we identify the range of contaminants present in commonly used DNA extraction reagents and demonstrate the significant impact they can have on microbiota studies.

## RESULTS

### 16S rRNA gene sequencing of a pure *Salmonella bongori* culture

To demonstrate the presence of contaminating DNA, and its impact on high and low biomass samples, we used 16S rRNA gene sequence profiling of a pure culture of *Salmonella bongori* that had undergone five rounds of serial ten-fold dilutions (equating to a range of approximately 10^8^ cells as input for DNA extraction in the original undiluted sample, to 10^3^ cells in dilution five). *S. bongori* was chosen because we have not observed it as a contaminant in any of our previous studies and it can be differentiated from other *Salmonella* species by sequencing. As a pure culture was used as starting template, regardless of starting biomass, any organisms other than *S. bongori* observed in subsequent DNA sequencing results must be derived from contamination. Aliquots from the dilution series were sent to three institutes (Imperial College London, ICL; University of Birmingham, UB; Wellcome Trust Sanger Institute, WTSI) and processed with different batches of the FastDNA Spin Kit for Soil (kit FP). 16S rRNA gene amplicons were generated using both 20 and 40 PCR cycles and returned to WTSI for Illumina MiSeq sequencing.

*S. bongori* was the sole organism identified in the original undiluted culture but with subsequent dilutions a range of contaminating bacterial groups increased in relative abundance while the proportion of *S. bongori* reads concurrently decreased (Fig. 1). By the fifth serial dilution, equivalent to an input biomass of roughly 10^3^ *Salmonella* cells, contamination was the dominant feature of the sequencing results. This pattern was consistent across all three sites and was most pronounced with 40 cycles of PCR. These results highlight a key problem with low biomass samples. The most diluted 20-PCR cycle samples resulted in low PCR product yields and so were under-represented in the sequencing mix. Conversely, using 40 PCR cycles generated enough PCR product for effective sequencing but a significant proportion of the resulting sequence data was derived from contaminating DNA. It should be noted though that even when using only 20 PCR cycles contamination was still predominant with the lowest input biomass (Supplementary Fig. SI).

**Figure 1.**
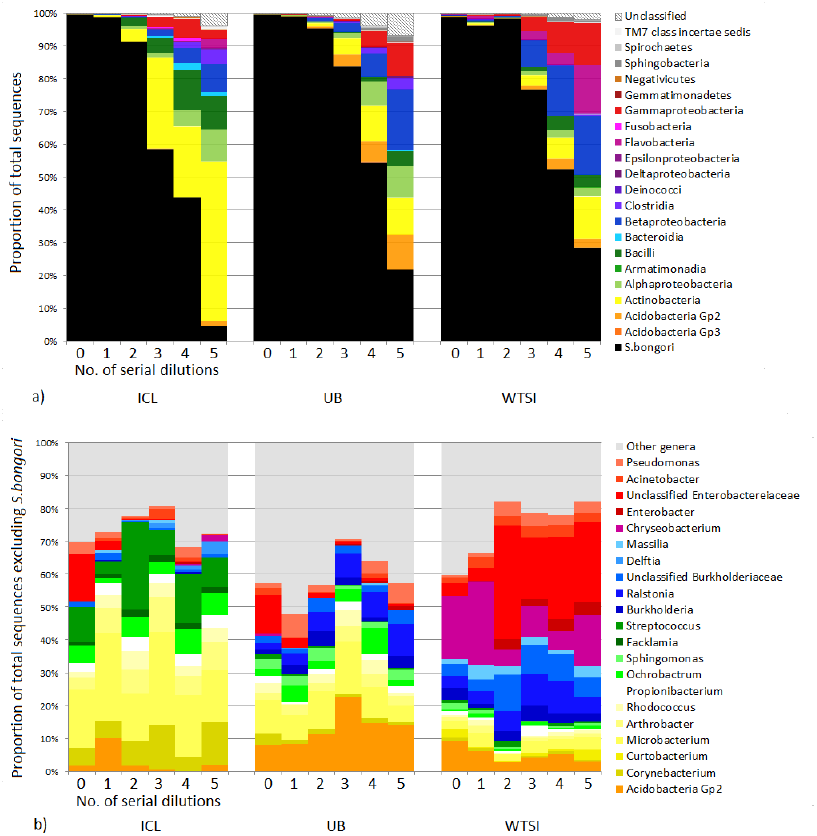
Summary of the 16S rRNA gene sequencing taxonomic assignment from ten-fold diluted pure cultures (undiluted DNA extractions contained approximately 10^8^ cells), extracted at ICL, UB and WTSI laboratories and amplified with 40 PCR cycles. a) Proportion of *S. bongori* sequence reads in black. Proportional abundance of non-*Salmonella* reads at the Class level are indicated by other colours. As the sample becomes more dilute, the proportion of the sequenced bacterial amplicons from the cultured microorganism decreases and contaminants become more dominant. b) Abundance of genera which make up >0.5% of the results from at least one laboratory, excluding *S. bongori*. The profiles of the non-*Salmonella* reads within each laboratory/kit batch are consistent but differ between sites.

Sequence profiles revealed some similar taxonomic classifications between all sites, including Acidobacteria Gp2, *Microbacterium, Propionibacterium,* and *Pseudomonas* (Fig. 1b). Differences between sites were observed however, with *Chryseobacterium, Enterobacter* and *Massilia* at WTSI, *Sphingomonas* at UB, and *Corynebacterium, Facklamia* and *Streptococcus* more dominant at ICL along with a greater proportion of Actinobacteria in general (Fig. 1a). This illustrates that there is variation in contaminant content between laboratories, and between reagent/kit batches. Many of the contaminating OTUs represent bacterial genera normally found in soil and water, for example *Arthrobacter, Burkholderia, Chryseobacterium, Ochrobactrum, Pseudomonas, Ralstonia, Rhodococcus* and *Sphingomonas,* while others, such as *Corynebacterium, Propionibacterium* and *Streptococcus,* are common human skin-associated organisms.

### Quantitative PCR of bacterial biomass

To assess how much background bacterial DNA was present in the samples, we performed qPCR of bacterial 16S rRNA genes and calculated the copy number of genes present with reference to a standard curve. In the absence of contamination copy number of the 16S rRNA genes present should correlate with dilution of *S. bongori* and reduce in a linear manner. However, at the third dilution copy number remained stable and did not reduce further, indicating the presence of background DNA at approximately 500 copies per µl of elution volume from the DNA extraction kit (Supplementary Fig. S2).

### Shotgun metagenomics of a pure *S. bongori* culture processed with four commercial DNA extraction kits

Having established that 16S rRNA gene sequencing results can be confounded by contaminating DNA we next investigated whether similar patterns emerge in shotgun metagenomics studies, which do not involve a targeted PCR step. We hypothesised that if contamination arises from the DNA extraction kit, it should also be present in metagenomic sequencing results. DNA extraction kits from four different manufacturers were used in order to investigate whether or not the problem was limited to a single manufacturer. Aliquots from the same *S. bongori* dilution series were processed at UB with the FastDNA Spin Kit For Soil (FP), MoBio UltraClean Microbial DNA Isolation Kit (MB), QIAmp DNA Stool Mini Kit (QIA) and PSP Spin Stool DNA Plus kit (PSP). As with 16S rRNA gene sequencing, it was found that as the sample biomass dilution increased, the proportion of reads mapping to the *S. bongori* reference genome sequence decreased (Fig. 2a). Regardless of kit, contamination was always the predominant feature of the sequence data by the fourth serial dilution, which equated to an input of around 10^4^ *Salmonella* cells.

**Figure 2.**
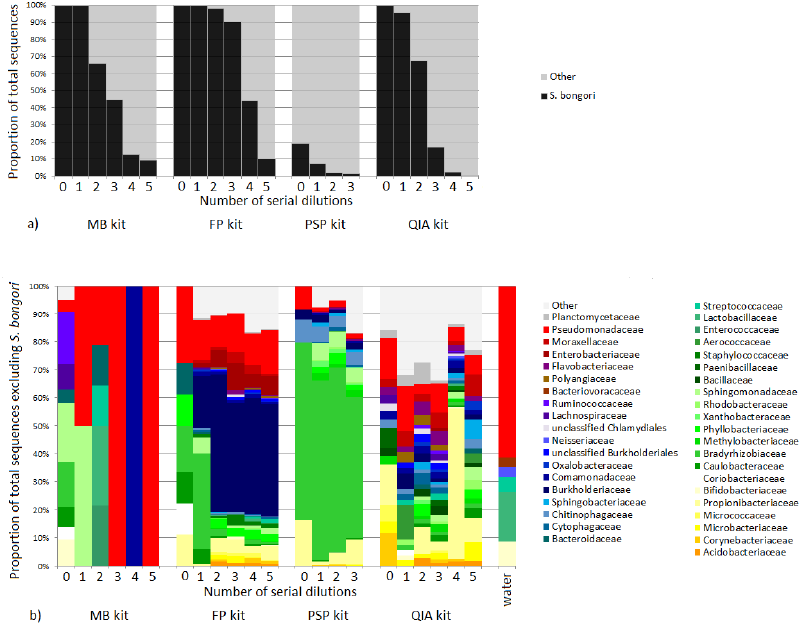
Summary of the metagenomic data for the *S. bongori* ten-fold dilution series (initial undiluted samples contained approximately 10^8^ cells), extracted with four different kits. a) As the starting material becomes more diluted, the proportion of sequenced reads mapping to the *S. bongori* reference genome decreases for all kits and contamination becomes more prominent. b) The profile of the non-*Salmonella* reads (grouped by Family, only those comprising >1% of reads from at least one kit are shown) is different for each of the four kits.

A range of environmental bacteria was observed, which were of a different profile in each kit (Fig. 2b). FP had a stable kit profile dominated by *Burkholderia,* PSP was dominated by *Bradyrhizobium,* while the QIA kit had the most complex mix of bacterial DNA. Bradyrhizobiaceae, Burkholderiaceae, Chitinophagaceae, Comomonadaceae, Propionibacteriaceae and Pseudomonadaceae were present in at least three quarters of the dilutions from PSP, FP and QIA kits. However, relative abundances of taxa at the Family level varied according to kit: FP was marked by Burkholderiaceae and Enterobacteriaceae, PSP was marked by Bradyrhizobiaceae and Chitinophagaceae. The contamination in the QIA kit was relatively diverse in comparison to the other kits, and included higher proportions of Aerococcaceae, Bacillaceae, Flavobacteriaceae, Microbacteriaceae, Paenibacillaceae, Planctomycetaceae and Polyangiaceae than the other kits. Kit MB did not have a distinct contaminant profile and varied from dilution to dilution due to paucity of reads.

These metagenomic results therefore clearly show that contamination becomes the dominant feature of sequence data from low biomass samples, and that the kit used to extract DNA can have an impact on the observed bacterial diversity, even in the absence of a PCR amplification step. Reducing input biomass again increases the impact of these contaminants upon the observed microbiota.

### Impact of contaminated extraction kits on a study of low-biomass microbiota

Having established that the contamination in different lots of DNA extraction kits is not constant or predictable, we next show the impact that this can have on real datasets. A recent study in a refugee camp on the border between Thailand and Burma used an existing nasopharyngeal swab archive^38^ to examine the development of the infant nasopharyngeal microbiota [unpublished]. A cohort of 20 children born in 2007-2008 were sampled every month until two years of age, and the 16S rRNA gene profiles of these swabs were sequenced by 454 pyrosequencing.

Principal coordinate analysis (PCoA)showed two distinct clusters distinguishing samples taken during early life from those taken from subsequent sampling time points, suggesting an early, founder nasopharyngeal microbiota (Fig. 3a). Four batches of FP kits had been used to extract the samples and a record was made of which kit was used for each sample. Further analysis of the OTUs present indicated that samples possessed different communities depending on which kit had been used for DNA extraction (Figs. 3b,3d, 3e) and that the first two kits’ associated OTUs made up the majority of their samples’ reads (Fig. 3d). As samples had been extracted in chronological order, rather than random order, this led to the false conclusion that OTUs from the first two kits were associated with age. OTUs driving clustering to the left in Figures 3a and 3b (*P* value of <0.01), were classified as *Achromobacter, Aminobacter, Brevundimonas, Herbaspirillum, Ochrobactrum, Pedobacter, Pseudomonas, Rhodococcus, Sphingomonas* and *Stenotrophomonas*. OTUs driving data points to the right *(P* value of <0.01), included *Acidaminococcus* and *Ralstonia*. A full list of significant OTUs is shown in Supplementary Table S1. Once the contaminants were identified and removed, the PCoA clustering of samples from the run no longer had a discernible pattern, showing that the contamination was the biggest driver of sample ordination (Fig. 3c). New aliquots were obtained from the original sample archive and were reprocessed using a different kit lot and sequenced [data not shown]. The previously observed contaminant OTUs were not detected, confirming their absence in the original nasopharyngeal samples.

**Figure 3.**
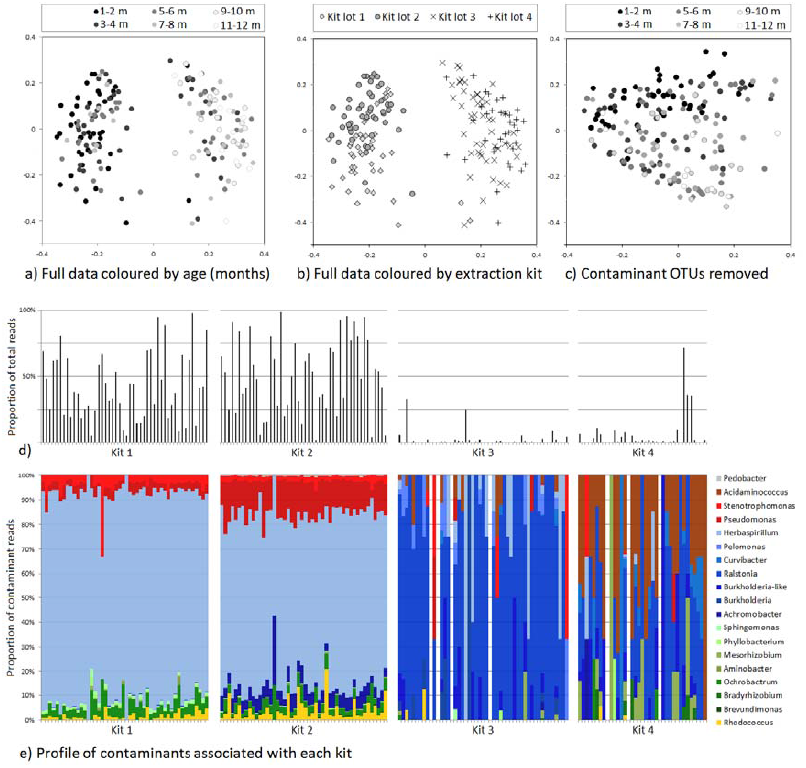
Summary of the contaminant content of nasopharyngeal samples from Thailand. a) PCoA plot appears to show age-related clustering, however, b) extraction kit lot explains the pattern better. c) When coloured by age, plot shows the loss of initial clustering pattern after excluding contaminant OTUs from ordination. d) The proportion of reads attributed to contaminant OTUs for each sample, demonstrating that the first two kits were the most heavily contaminated. e) Genus-level profile of contaminant OTUs for each kit used.

This dataset therefore serves as a case study for the significant, and potentially misleading, impact that contaminants originating from kits can have on microbiota analyses and subsequent conclusions.

## DISCUSSION

Results presented here show that contamination with bacterial DNA or cells in DNA extraction kit reagents should not only be a concern for 16S rRNA gene sequencing projects, which require PCR amplification, but also for shotgun metagenomics projects.

Contaminating DNA has been reported from PCR reagents, kits and water many times^3,15,17^. The taxa identified are mostly soil- or water-dwelling bacteria and are frequently associated with nitrogen fixation. One explanation for this may be that nitrogen is often used instead of air in ultrapure water storage tanks^3^. Contamination of DNA extraction kit reagents has also been reported^16^ and kit contamination is a particular challenge for low biomass studies, which may provide little template DNA to compete with that in the reagents for amplification^12,39^. Issues of contamination have plagued studies, with high-profile examples in the fields of novel virus discovery, such as in the false association of XMRV and chronic fatigue syndrome^40^, and the study of ancient DNA of early humans and pathogens^41,42^. The microbial content of ancient ice core samples has also shown to be inconsistent when analysed by different laboratories^39^.

The importance of this issue when analysing low biomass samples, despite such high-profile reports of reagent contamination, apparently remains underappreciated in the microbiota research community. Well-controlled studies, such as in Segal *et al*. who examined the lung microbiota through bronchoalveolar lavage sampling, report their results against the background of copious sequenced ‘background’ controls^43^. However, many recent DNA sequence-based publications that describe the microbial communities of low-biomass environments do not report DNA quantification on initial samples, sequencing of negative controls or describe their contaminant removal or identification procedures. Our literature searches have indicated that there are a number of low biomass microbiota studies that report taxa, often statistically noteworthy or core members, that overlap with those we report here from our negative control kit reagents and water (shown in Table 1). While it is possible that the suspect taxa are genuinely present in these samples, in many cases they are biologically unexpected: for example, rhizosphere-associated bacteria that have been implicated in human disease^27,44^. Tellingly, Laurence *et al*.^18^ recently demonstrated with an *in silico* analysis that *Bradyrhizobium* is a common contaminant of sequencing datasets including the 1000 Human Genome Project. Having demonstrated the critical impact that contaminating DNA may have on conclusions drawn from sequence-based data, it becomes important to be able to determine which observations are genuine.

A number of methods have been devised to treat reagents in order to reduce potential contamination, including: gamma^45^ or UV radiation^13,46–48^; DNase treatment^10,13,47,49–51^ restriction digests^10,13,47,52,53^ caesium chloride density gradient centrifugation^10^, and DNA intercalation and crosslinking with 8-methoxypsoralen^47,54^ propidium monoazide^55^ or ethidium monoazide^56,57^. However tests of these methods show varying levels of success. Radiation may reduce the activity of enzymes, DNase inactivation can also damage the polymerase, restriction enzymes may introduce more contaminating DNA, and unbound DNA intercalators inhibit amplification of the intended template^56,58^. An alternative to decontamination is to preferentially amplify the template DNA using broad range primer extension PCR^59^ but this, and the treatment of the PCR reagents, cannot account for contamination introduced through DNA extraction kits. An *in silico* approach for microbiota studies is to identify contaminants that are sequenced using negative controls or contaminant databases in order to screen them out of downstream analysis^17,60^.

By adding negative sequencing controls (specifically, template-free “blanks” processed with the same DNA extraction and PCR amplification kits as the real samples, sequenced on the same run) it is possible to identify reads originating from contamination, and distinguish them from those derived from actual constituent taxa. We have developed a set of recommendations that may help to limit the impact of reagent contamination (Box 1). With awareness of common contaminating species, careful collection of controls to cover different batches of sampling, extraction and PCR kits, and sequencing to monitor the content of these controls, it should be possible to effectively mitigate the impact of contaminants in microbiota studies.

## CONCLUSION

We have shown that bacterial DNA contamination in extraction kits and laboratory reagents can significantly influence the results of microbiota studies, particularly when investigating samples containing a low microbial biomass. Such contamination is a concern for both 16S rRNA gene sequencing projects, which require targeted PCR amplification and enrichment, but also for shotgun metagenomic projects which do not. Awareness of this issue by the microbiota research community is important to ensure that studies are adequately controlled and erroneous conclusions are not drawn from culture-independent investigations.

### BOX 1

Recommendations to reduce the impact of contaminants in sequence-based, low-biomass microbiota studies.

1. Maximise the starting sample biomass by choice of sample type. If microbial load is less than approximately 10^3^ to 10^4^ cells it may not be possible to obtain robust results as contamination appears to predominate.
2. Minimise risk of contamination at the point of sample collection. PCR and extraction kit reagents may be treated to reduce contaminant DNA.
3. Collect process and sequence technical controls from each batch of sample collection/storage medium, each extraction kit, and each PCR kit concurrently with the environmental samples of interest.
4. Samples should be processed in random order to avoid creating false patterns and ideally carried out in replicates, which should be processed using different kit/reagent batches.
5. A record should be made of which sample was processed with which kit so that contamination of a particular kit lot number can be traced through to the final dataset.
6. Quantification of the negative controls and samples should be ongoing during processing in order to monitor contamination as it arises.
7. After sequencing, be wary of taxa that are present in the negative controls, taxa that are statistically associated with a particular batch of reagents, and taxa that are unexpected biologically and also coincide with previously reported contaminants such as those listed in Table 1.
8. In the event that suspect taxa are still of interest, repeat sequencing should be carried out on additional samples using separate batches of DNA extraction kits/reagents, and ideally a non-sequencing based approach (such as Fluorescent *in situ* hybridisation, using properly validated probe sets) should also be used to further confirm their presence in the samples.

## METHODS

### Samples

For the 16S rRNA gene and metagenomic profiling, *Salmonella bongori* strain NCTC-12419 was cultured overnight on LB plates without antibiotics, at 37 °C. A single colony was used to inoculate an LB broth, which was incubated with shaking at 37 °C overnight. The OD_600_ upon retrieval was 1.62, equating to around 10^9^ CFU/ml. 20 µl from the culture was plated out on LB and observed to be a pure culture after overnight incubation. Five ten-fold dilutions from the starter culture were made in fresh LB. 1 ml aliquots of each dilution were immediately stored at –80 °C, and duplicates shipped on dry ice to Imperial College London and the University of Birmingham.

For the nasopharyngeal microbiota study, the samples were nasopharyngeal swabs collected from a cohort of infants in Maela refugee camp in Thailand as described previously^38^. These were vortexed in STGG medium and then stored at –80 °C.

### DNA extraction

For the 16S rRNA gene profiling work, each of the three institutes (Imperial College London, ICL; University of Birmingham, UB; Wellcome Trust Sanger Institute, WTSI) extracted DNA from the *S. bongori* aliquots in parallel, using different production batches of the FastDNA Spin Kit For Soil (MP Biomedicals, kit lots #38098, #15447 and #30252), according to the manufacturer’s protocol. UB and WTSI extracted DNA from 200 µl of sample and eluted in 50 µl; ICL extracted from 500 µl of sample and eluted in 100 µl. This meant that our DNA extractions across the five-fold serial dilutions spanned a range of sample biomass from approximately 10^8^ down to 10^3^ cells.

For the metagenomic sequencing, 200 µl aliquots of each *S. bongori* dilution and negative controls were processed using four commercially available DNA extraction kits at UB. The final elution volume for all kits was 100 µl per sample. The FP kit (lot #38098) was used according to the manufacturer’s protocol, with the exception of the homogeniser step. This was performed with a Qiagen Tissue Lyser: 1 minute at speed 30/s followed by 30 seconds cooling the tubes on ice, repeated 3 times. The UltraClean Microbial DNA Isolation Kit (MO BIO Laboratories) (kit MB, lot #U13F22) was used according to the manufacturer’s protocol with the exception of homogenisation, which was replaced by 10 minutes of vortexing. The QIAmp DNA Stool Mini Kit (Qiagen) (kit QIA, lot #145036714) was used according to the manufacturer’s stool pathogen detection protocol. The heating step was at 90 °C. The PSP Spin Stool DNA Plus kit (STRATEC Molecular) (kit PSP, lot #JB110047) was used according to the manufacturer’s stool homogenate protocol.

For the nasopharyngeal microbiota study, a 200 µl aliquot was taken from each sample and processed with the manufacturer’s vortex modification of the FP kit protocol. DNA was then shipped to WTSI for further processing and sequencing (see below).

### qPCR

A standard curve was produced by cloning the near full-length 16S rRNA gene of *Vibrio natriegens* DSMZ 759 amplified using primers 27F and 1492R^61^ into the TOPO TA vector (Life Technologies), quantifying using fluorescent assay (Quant-IT, Life Technologies) and diluting to produce a standard curve from 10^8^ to 10^3^ copies per µl. A ViiA 7 Real-time PCR system (Life Technologies) with KAPA Biosytems SYBR Fast qPCR Master Mix was used to perform quantitative PCR of the V4 region of the bacterial 16S rRNA gene for each *S. bongori* dilution extraction. Primers used were: S-D-Bact-0564-a-S-15, 5’-AYTGGGYDTAAAGNG and S-D-Bact-0785-b-A-18, 5-TACNVGGGTATCTAATCC^62^ generating a 253 bp amplicon. 15 µl reactions were performed in triplicate and included template-free controls. Reactions consisted of 0.3 µl of 10 µM dilutions of each primer, 7.5 µl of SYBR Fast mastermix and 1.9 µl of microbial DNA free PCR water (MOBIO) and 5 µl of 1:5 diluted template (to avoid pipetting less than 5 µl). Cycle conditions were 90 °C for 3 minutes followed by 40 cycles of: 95 °C for 20 seconds; 50 °C for 30 seconds; and 72 °C for 30 seconds. Melt curves were run from 60 to 95 °C over 15 minutes.

### Sequencing

Samples for the *S. bongori* culture 16S rRNA gene profiling were PCR-amplified using barcoded fusion primers targeting the V1-V2 region of the gene (27f_Miseq: AATGATACGGCGACCACCGAGATCTACAC TATGGTAATT CC AGMGTTYGATYMTGGCTCAG and 338R_MiSeq: CAAGCAGAAGACGGCATACGAGAT nnnnnnnnnnnn AGTCAGTCAG AA GCTGCCTCCCGTAGGAGT, where the n string represents unique 12-mer barcodes used for each sample studied) and then sequenced on the Illumina MiSeq platform using 2 × 250 bp cycles. The PCR amplification was carried out with Q5 (New England Biolabs) at WTSI, ICL and UB, using fresh reagents and consumables, autoclaved microcentrifuge tubes, filtered pipette tips, and performed in a hood to reduce the risk of airborne contamination. Each sample was amplified with both 20 and 40 PCR cycles under the following conditions: 94 °C for 30 seconds, 53 °C for 30 seconds, 68 °C for 2 minutes. Negative controls were included for each batch.

For metagenomic sequencing, all samples were quantified using Nanodrop (Thermo Scientific) and Qubit (Life Technologies) machines, and did not need to be diluted before Illumina Nextera XT library preparation (processed according to manufacturer’s protocol). Libraries were multiplexed on the Illumina MiSeq in paired 250-base mode following a standard MiSeq wash protocol.

For the nasopharyngeal microbiota study, DNA extractions from 182 swabs were PCR-amplified and barcoded for sequencing the 16S rRNA gene V3-V5 region on the 454 platform as described previously^63^.

### Sequence analysis

For the 16S rRNA gene profiling, data was processed using mothur^64^. The mothur MiSeq SOP^65^ was followed with the exception of screen.seqs, which used the maximum length of the 97.5 percentile value, and chimera checking, which was performed with Perseus^66^ instead of UCHIME.

For the metagenomic profiling, reads were quality checked and trimmed for low-quality regions and adaptor sequences using Trimmomatic^67^. Similarity sequencing for taxonomic assignments was performed using LAST in 6-frame translation mode against the Microbial RefSeq protein database^68^. Taxonomic assignments were determined with MEGAN, which employs a lowest common ancestor (LCA) to taxonomic assignments, using settings Min Support 2, Min Score 250, Max Expected 0.1, Top Percent 10.0^69^.

For the nasopharyngeal microbiota study, the data were processed, cleaned and analysed using the mothur Schloss SOP^70^ and randomly subsampled to 200 sequence reads per sample. As part of the contamination identification procedure, the metastats package^71^ within mothur was used to identify OTUs that were significantly associated with each extraction kit batch. Jaccard PCoA plots were generated with mothur, comparing the dataset with and without these flagged OTUs included.

## ACKNOWLEDGEMENTS

SJS, JP, AWW and sequencing costs were supported by the Wellcome Trust [grant number 098051]. MJC was supported by a Wellcome Trust Centre for Respiratory Infection Basic Science Fellowship. STC is funded by the National Institute for Health Research (NIHR). WOC and MFM are supported by a Wellcome Trust Joint Senior Investigator’s Award, which also supports EMT. PT was supported by a Wellcome Trust Clinical Training Fellowship [grant number 083735/Z/07/Z], NJL is supported by a Medical Research Council Special Training Fellowship in Biomedical Informatics. The views expressed are those of the authors and not necessarily those of the Wellcome Trust, the NHS, the NIHR, or the Department of Health.

We would like to thank the Wellcome Trust Sanger Institute’s core sequencing team, Paul Scott for his assistance in the laboratory and for providing a list of contaminants derived from multiple displacement amplification kits, and Phil James for assistance with qPCR.

### AUTHOR CONTRIBUTIONS

SJS, MJC, NL and AWW devised experiments. SJS, ET, STC and NL performed experiments. SJS, MJC, NL and AWW analysed data and prepared figures. PT provided data demonstrating the impact of extraction kit contaminants. WOC, MFM, NL and JP provided resources, guidance and support. SJS, MJC, NL and AWW wrote the paper. All authors read and approved the final manuscript.

### COMPETING FINANCIAL INTERESTS

The authors have no competing financial interests.

## References

1. Kunin, V., Engelbrektson, A., Ochman, H. & Hugenholtz, P. Wrinkles in the rare biosphere: pyrosequencing errors can lead to artificial inflation of diversity estimates. Environmental Microbiology 12,118–123 (2010).

2. Wintzingerode, F. v., Göbel, U. B. & Stackebrandt, E. Determination of microbial diversity in environmental samples: pitfalls of PCR-based rRNA analysis. FEMS Microbiology Reviews 21,213–229 (1997).

3. Kulakov, L. A., McAlister, M. B., Ogden, K. L., Larkin, M. J. & O’Hanlon, J. F. Analysis of bacteria contaminating ultrapure water in industrial systems. Applied and Environmental Microbiology 68, 1548–1555 (2002).

4. McAlister, M. B., Kulakov, L. A., O’Hanlon, J. F., Larkin, M. J. & Ogden, K. L. Survival and nutritional requirements of three bacteria isolated from ultrapure water. Journal of Industrial Microbiology and Biotechnology 29, 75–82 (2002).

5. Kéki, Z., Grébner, K., Bohus, V., Márialigeti, K. & Tóth, E. M. Application of special oligotrophic media for cultivation of bacterial communities originated from ultrapure water. Acta Microbiologica et Immunologica Hungarica 60, 345–357 (2013).

6. Bohus, V. et al. Bacterial communities in an ultrapure water containing storage tank of a power plant. Acta Microbiologica et Immunologica Hungarica 58, 371–382 (2011).

7. McFeters, G. A., Broadaway, S. C., Pyle, B. H. & Egozy, Y. Distribution of bacteria within operating laboratory water purification systems. Applied and Environmental Microbiology 59, 1410–1415 (1993).

8. Nogami, T., Ohto, T., Kawaguchi, O., Zaitsu, Y. & Sasaki, S. Estimation of bacterial contamination in ultrapure water: application of the anti-DNA antibody. Analytical Chemistry 70, 5296–5301 (1998).

9. Shen, H., Rogelj, S. & Kieft, T. L. Sensitive, real-time PCR detects low-levels of contamination by Legionella pneumophila in commercial reagents. Molecular and Cellular Probes 20,147–153 (2006).

10. Rand, K. H. & Houck, H. *Taq* polymerase contains bacterial DNA of unknown origin. Molecular and Cellular Probes 4, 445–450 (1990).

11. Maiwald, M., Ditton, H. J., Sonntag, H. G. & von Knebel Doeberitz, M. Characterization of contaminating DNA in *Taq* polymerase which occurs during amplification with a primer set for Legionella 5S ribosomal RNA. Molecular and Cellular Probes 8,11–14 (1994).

12. Tanner, M. A., Goebel, B. M., Dojka, M. A. & Pace, N. R. Specific ribosomal DNA sequences from diverse environmental settings correlate with experimental contaminants. Applied and Environmental Microbiology 64, 3110–3113 (1998).

13. Corless, C. E. et al. Contamination and sensitivity issues with a real-time universal 16S rRNA PCR. Journal of Clinical Microbiology 38,1747–1752 (2000).

14. Grahn, N., Olofsson, M., Ellnebo-Svedlund, K., Monstein, H. J. & Jonasson, J. Identification of mixed bacterial DNA contamination in broad-range PCR amplification of 16S rDNA V1 and V3 variable regions by pyrosequencing of cloned amplicons. FEMS Microbiology Letters 219, 87–91 (2003).

15. Newsome, T., Li, B. J., Zou, N. & Lo, S. C. Presence of bacterial phage-like DNA sequences in commercial Taq DNA polymerase reagents. Journal of Clinical Microbiology 42, 2264–2267 (2004).

16. Mohammadi, T., Reesink, H. W., Vandenbroucke-Grauls, C. M. & Savelkoul, P. H. Removal of contaminating DNA from commercial nucleic acid extraction kit reagents. Journal of Microbiological Methods 61, 285–288 (2005).

17. Barton, H. A., Taylor, N. M., Lubbers, B. R. & Pemberton, A. C. DNA extraction from low-biomass carbonate rock: an improved method with reduced contamination and the low-biomass contaminant database. Journal of Microbiological Methods 66, 21–31 (2006).

18. Laurence, M., Hatzis, C. & Brash, D. E. Common contaminants in next-generation sequencing that hinder discovery of low-abundance microbes. PLoS One 9, e97876 (2014).

19. Oberauner, L. et al. The ignored diversity: complex bacterial communities in intensive care units revealed by 16S pyrosequencing. Scientific Reports 3,1413 (2013).

20. La Duc, M. T., Kern, R. & Venkateswaran, K. Microbial monitoring of spacecraft and associated environments. Microbial Ecology 47,150–158 (2004).

21. Ling, Z. et al. Pyrosequencing analysis of the human microbiota of healthy Chinese undergraduates. BMC Genomics 14, 390 (2013).

22. Benítez-Páez, A. et al. Detection of transient bacteraemia following dental extractions by 16S rDNA pyrosequencing: a pilot study. PLoS One 8, e57782 (2013).

23. Amar, J. et al. Involvement of tissue bacteria in the onset of diabetes in humans: evidence for a concept. Diabetologia 54, 3055–3061 (2011).

24. Branton, W. G. et al. Brain microbial populations in HIV/AIDS: α-proteobacteria predominate independent of host immune status. PLoS One 8, e54673 (2013).

25. Borewicz, K. et al. Longitudinal analysis of the lung microbiome in lung transplantation. FEMS Microbiology Letters 339, 57–65 (2013).

26. Dong, Q. et al. Diversity of bacteria at healthy human conjunctiva. Investigative Ophthalmology and Visual Science 52, 5408–5413 (2011).

27. Xuan, C. et al. Microbial dysbiosis is associated with human breast cancer. PLoS One 9, e83744 (2014).

28. Kuehn, J. S. et al. Bacterial community profiling of milk samples as a means to understand culture-negative bovine clinical mastitis. PLoS One 8, e61959 (2013).

29. Srinivas, G. et al. Genome-wide mapping of gene-microbiota interactions in susceptibility to autoimmune skin blistering. Nature Communications 4, 2462 (2013).

30. Boissière, A. et al. Midgut microbiota of the malaria mosquito vector *Anopheles gambiae* and interactions with *Plasmodium falciparum* infection. PLoS Pathogens 8, e1002742 (2012).

31. McKenzie, V. J., Bowers, R. M., Fierer, N., Knight, R. & Lauber, C. L. Co-habiting amphibian species harbor unique skin bacterial communities in wild populations. ISME Journal 6, 588–596 (2012).

32. Carlos, C., Torres, T. T. & Ottoboni, L. M. Bacterial communities and species-specific associations with the mucus of Brazilian coral species. Scientific Reports 3,1624 (2013).

33. Cheng, X. Y. et al. Metagenomic analysis of the pinewood nematode microbiome reveals a symbiotic relationship critical for xenobiotics degradation. Scientific Reports 3,1869 (2013).

34. Davidson, S. K., Powell, R. & James, S. A global survey of the bacteria within earthworm nephridia. Molecular Phylogenetics and Evolution 67,188–200 (2013).

35. Knowlton, C., Veerapaneni, R., D’Elia, T. & Rogers, S. O. Microbial analyses of ancient ice core sections from Greenland and Antarctica. Biology 2, 206–232 (2013).

36. Shtarkman, Y. M. et al. Subglacial Lake Vostok (Antarctica) accretion ice contains a diverse set of sequences from aquatic, marine and sediment-inhabiting bacteria and eukarya. PLoS One 8, e67221 (2013).

37. DeLeon-Rodriguez, N. et al. Microbiome of the upper troposphere: Species composition and prevalence, effects of tropical storms, and atmospheric implications. Proceedings of the National Academy of Sciences of the United States of America 110, 2575–2580 (2013).

38. Turner, P. et al. A longitudinal study of *Streptococcus pneumoniae* carriage in a cohort of infants and their mothers on the Thailand-Myanmar border. PLoS One 7, e38271 (2012).

39. Willerslev, E., Hansen, A. J. & Poinar, H. N. Isolation of nucleic acids and cultures from fossil ice and permafrost. Trends in Ecology and Evolution 19,141–147 (2004).

40. Kearney, M. F. et al. Multiple sources of contamination in samples from patients reported to have XMRV infection. PLoS One 7, e30889 (2012).

41. Cooper, A. & Poinar, H. N. Ancient DNA: Do It Right or Not at All. Science 289,1139 (2000).

42. Roberts, C. & Ingham, S. Using ancient DNA analysis in palaeopathology: a critical analysis of published papers, with recommendations for future work. International Journal of Osteoarchaeology 18, 600–613 (2008).

43. Segal, L. N. et al. Enrichment of lung microbiome with supraglottic taxa is associated with increased pulmonary inflammation. Microbiome 1,19 (2013).

44. Bhatt, A. S. et al. Sequence-based discovery of Bradyrhizobium enterica in cord colitis syndrome. New England Journal of Medicine 369, 517–528 (2013).

45. Deragon, J.-M., Sinnett, D., Mitchell, G., Potier, M. & Labuda, D. Use of gamma irradiation to eliminate DNA contamination for PCR. Nucleic Acids Research 18, 6149 (1990).

46. Sarkar, G. & Sommer, S. S. Shedding light on PCR contamination. Nature 343, 27 (1990).

47. Klaschik, S., Lehmann, L. E., Raadts, A., Hoeft, A. & Stuber, F. Comparison of different decontamination methods for reagents to detect low concentrations of bacterial 16S DNA by real-time-PCR. Molecular Biotechnology 22, 231–242 (2002).

48. Tamariz, J., Voynarovska, K., Prinz, M. & Caragine, T. The application of ultraviolet irradiation to exogenous sources of DNA in plasticware and water for the amplification of low copy number DNA. Journal of Forensic Sciences 51, 790–794 (2006).

49. Hilali, F., Saulnier, P., Chachaty, E. & Andremont, A. Decontamination of polymerase chain reaction reagents for detection of low concentrations of 16S rRNA genes. Molecular Biotechnology 3, 207–216 (1997).

50. Heininger, A. et al. DNase pretreatment of master mix reagents improves the validity of universal 16S rRNA gene PCR results. Journal of Clinical Microbiology 41, 1763–1765 (2003).

51. Silkie, S. S., Tolcher, M. P. & Nelson, K. L. Reagent decontamination to eliminate false-positives in *Escherichia coli* q PCR. Journal of Microbiological Methods 72, 275–282 (2008).

52. Carroll, N. M., Adamson, P. & Okhravi, N. Elimination of bacterial DNA from Taq DNA polymerases by restriction endonuclease digestion. Journal of Clinical Microbiology 37, 3402–3404 (1999).

53. Mohammadi, T., Reesink, H. W., Vandenbroucke-Grauls, C. M. & Savelkoul, P. H. Optimization of real-time PCR assay for rapid and sensitive detection of eubacterial 16S ribosomal DNA in platelet concentrates. Journal of Clinical Microbiology 41, 4796–4798 (2003).

54. Hughes, M. S., Beck, L.-A. & Skuce, R. A. Identification and elimination of DNA sequences in Taq DNA polymerase. Journal of Clinical Microbiology 32, 2007–2008 (1994).

55. Vaishampayan, P. et al. New perspectives on viable microbial communities in low-biomass cleanroom environments. ISME Journal 7, 312–324 (2013).

56. Rueckert, A. & Morgan, H. W. Removal of contaminating DNA from polymerase chain reaction using ethidium monoazide. Journal of Microbiological Methods 68, 596–600 (2007).

57. Patel, P. et al. Development of an ethidium monoazide-enhanced internally controlled universal 16S rDNA real-time polymerase chain reaction assay for detection of bacterial contamination in platelet concentrates. Transfusion 52, 1423–1432 (2012).

58. Champlot, S. B. C, Pruvost, M., Bennett, E. A., Grange, T. & Geigl, E. M. An efficient multistrategy DNA decontamination procedure of PCR reagents for hypersensitive PCR applications. PLoS One 5, e13042 (2010).

59. Chang, S. S., Hsu, H. L., Cheng, J. C. & Tseng, C. P. An efficient strategy for broad-range detection of low abundance bacteria without DNA decontamination of PCR reagents. PLoS One 6, e20303 (2011).

60. Bonfert, T., Csaba, G., Zimmer, R. & Friedel, C. C. Mining RNA-Seq Data for Infections and Contaminations. PLoS One 8, e73071 (2013).

61. Lane, D. J. in Nucleic Acid Techniques in Bacterial Systematics (eds E; Stackebrandt & M; Goodfellow) 115–175 (Wiley, 1991).

62. Klindworth, A. et al. Evaluation of general 16S ribosomal RNA gene PCR primers for classical and next-generation sequencing-based diversity studies. Nucleic Acids Research 41, e1 (2013).

63. Cooper, P. et al. Patent human infections with the whipworm, Trichuris trichiura, are not associated with alterations in the faecal microbiota. PLoS One 8, e76573 (2013).

64. Schloss, P. D. et al. Introducing mothur: open-source, platform-independent, community-supported software for describing and comparing microbial communities. Applied and Environmental Microbiology 75, 7537–7541 (2009).

65. Kozich, J. J., Westcott, S. L., Baxter, N. T., Highlander, S. K. & Schloss, P. D. Development of a dual-index sequencing strategy and curation pipeline for analyzing amplicon sequence data on the MiSeq Illumina sequencing platform. Applied and Environmental Microbiology 79, 5112–5120 (2013).

66. Quince, C., Lanzen, A., Davenport, R. J. & Turnbaugh, P. J. Removing noise from pyrosequenced amplicons. BMC Bioinformatics 12, 38 (2011).

67. Bolger, A. M., Lohse, M. & Usadel, B. Trimmomatic: a flexible trimmer for Illumina sequence data. Bioinformatics [IN PRESS] (2014).

68. Kiełbasa, S. M., Wan, R., Sato, K., Horton, P. & Frith, M. C. Adaptive seeds tame genomic sequence comparison. Genome Research 21, 487–493 (2011).

69. Huson, D. H., Mitra, S., Ruscheweyh, H. J., Weber, N. & Schuster, S. C. Integrative analysis of environmental sequences using MEGAN4. Genome Research 21,1552–1560 (2011).

70. Schloss, P. D., Gevers, D. & Westcott, S. L. Reducing the effects of PCR amplification and sequencing artifacts on 16S rRNA-based studies. PLoS One 6, e27310 (2011).

71. White, J. R., Nagarajan, N. & Pop, M. Statistical methods for detecting differentially abundant features in clinical metagenomic samples. PLoS Computational Biology 5, e1000352 (2009).

