## Supplemental Figs 1 &#x26; 2; Supplemental Table 1 for "Reagent contamination can critically impact sequence-based microbiome analyses"

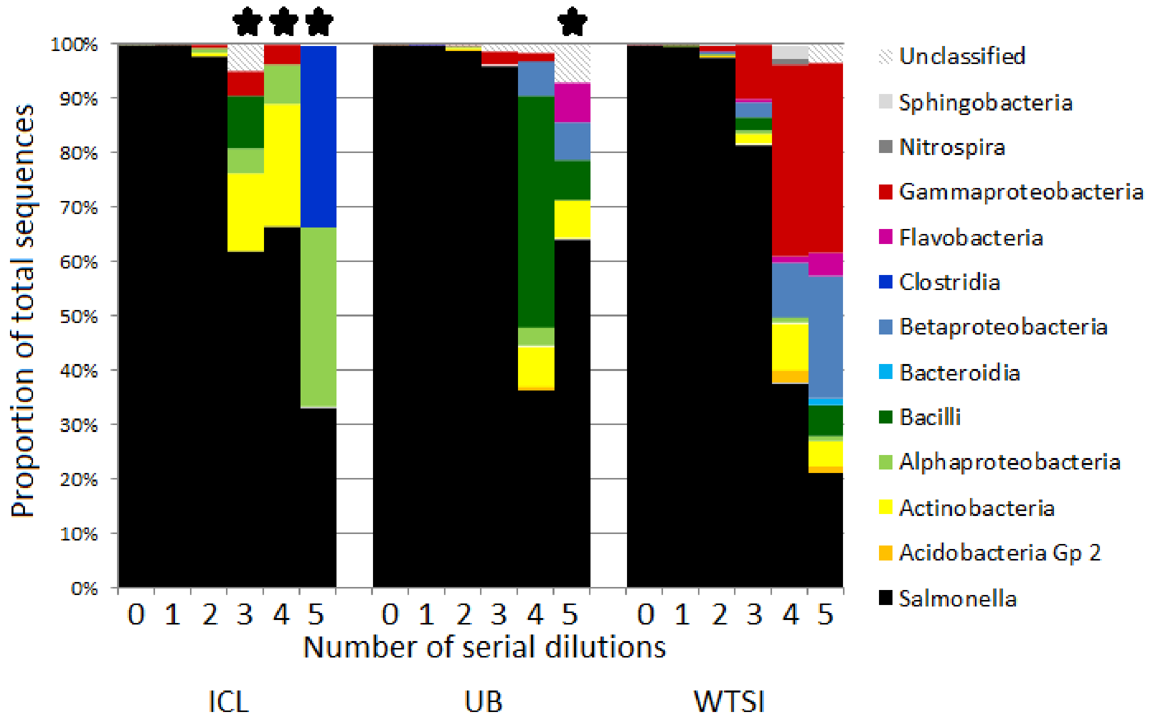


**Supplementary Figure S1:**

**16S rRNA gene profile of *S. bongori* pure culture serial dilutions amplified with 20 PCR cycles.**

*S. bongori* is shown in black; other taxa are grouped by Class. Stars mark samples that had fewer than 50 sequence reads: the most diluted samples gave no visible bands on electrophoresis gels after 20 PCR cycles, and these samples therefore tended to be under-represented in the sequence libraries. The centres that performed the DNA extraction and PCR steps are shown at the bottom of the figure (ICL = Imperial College London, UB = University of Birmingham, WTSI = Wellcome Trust Sanger Institute).

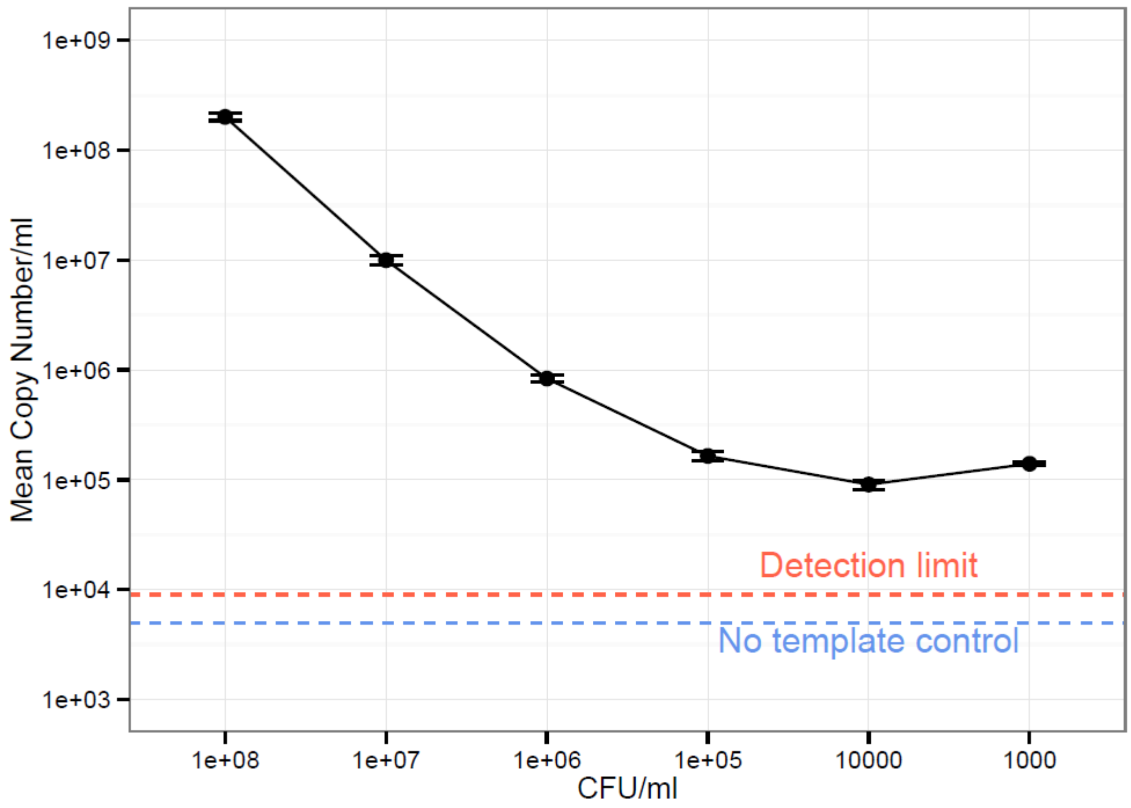


**Supplementary Figure S2:**

**Copy number of total 16S rRNA genes present in a dilution series of *S. bongori* culture.**

Total bacterial DNA present in serial ten-fold dilutions of a pure *S. bongori* culture was quantified using qPCR. While the copy number initially reduces in tandem with increased dilution, plateauing after four dilutions indicates consistent background levels of contaminating DNA. Error bars indicate standard deviation of triplicate reactions. The broken red line indicates the detection limit of 45 copies of 16S rRNA genes. The no template internal control for the qPCR reactions (shown in blue) was below the cycle threshold selected for interpreting the fluorescence values (i.e. less than 0), indicating the contamination did not come from the qPCR reagents themselves.

**Supplementary Table S1:**

**OTUs with significant correlation in PCoA plot Figures 3b and 3c.**

All taxa with a p-value of <0.05 are shown, with *P*<0.01 highlighted in bold. Although the data is from human nasopharyngeal swabs, many of the taxa are environmental bacteria associated with the DNA extraction kit.

| OTU identity | Classification | Environmental/human associated | X axis | | Y axis | |
|---|---|---|---|---|---|---|
| | | | Correlation coefficient | p-value | Correlation coefficient | p-value |
| **Otu003** | *Herbaspirillum* | Environmental | -0.753 | **0.000** | | |
| **Otu009** | *Pseudomonas* | Both | -0.591 | **0.000** | 0.250 | **0.001** |
| **Otu012** | *Ochrobactrum* | Environmental | -0.568 | **0.000** | | |
| **Otu014** | *Rhodococcus* | Environmental | -0.486 | **0.000** | 0.198 | **0.007** |
| **Otu030** | *Pedobacter* | Environmental | -0.438 | **0.000** | 0.264 | **0.000** |
| **Otu040** | *Aminobacter* | Environmental | -0.435 | **0.000** | | |
| **Otu025** | *Sphingomonas* | Environmental | -0.403 | **0.000** | -0.159 | 0.030 |
| **Otu031** | *Brevundimonas* | Environmental | -0.374 | **0.000** | 0.249 | **0.001** |
| **Otu015** | *Stenotrophomonas* | Both | -0.366 | **0.000** | -0.163 | 0.027 |
| **Otu013** | *Achromobacter* | Environmental | -0.357 | **0.000** | | |
| **Otu026** | *Phyllobacterium* | Environmental | -0.277 | **0.000** | | |
| **Otu116** | *Afipia* | Environmental | -0.208 | **0.005** | | |
| **Otu092** | *Moraxella* | Human | -0.152 | 0.040 | | |
| **Otu004** | *Haemophilus* | Human | 0.151 | 0.041 | | |
| **Otu081** | *Pseudonocardia* | Environmental | 0.151 | 0.041 | 0.202 | **0.006** |
| **Otu067** | *Bradyrhizobium* | Environmental | 0.154 | 0.037 | | |
| **Otu007** | *Corynebacterium* | Human | 0.157 | 0.033 | | |
| **Otu036** | *Burkholderia* | Both | 0.159 | 0.031 | | |
| **Otu060** | *Curvibacter* | Environmental | 0.164 | 0.026 | | |
| **Otu016** | *Ralstonia* | Environmental | 0.195 | **0.008** | 0.148 | 0.044 |
| **Otu017** | *Acidaminococcus* | Environmental | 0.210 | **0.004** | | |
| **Otu006** | *Moraxella* | Human | 0.232 | **0.001** | | |
| **Otu008** | Unclassified *Flavobacteriaceae* | Human | 0.242 | **0.001** | | |
| **Otu010** | *Helcococcus* | Human | 0.380 | **0.000** | | |
| **Otu001** | *Moraxella* | Human | 0.404 | **0.000** | | |
| **Otu197** | *Bordetella* | Both | | | 0.147 | 0.046 |
| **Otu090** | *Aeromonas* | Environmental | | | 0.148 | 0.045 |
| **Otu161** | *Kineosphaera* | Environmental | | | 0.150 | 0.042 |
| **Otu250** | *Perlucidibaca* | Environmental | | | 0.153 | 0.038 |
| **Otu117** | *Rheinheimera* | Environmental | | | 0.153 | 0.038 |
| **Otu058** | *Dyella* | Environmental | | | 0.153 | 0.038 |
| **Otu020** | *Actinobacillus* | Human | | | 0.154 | 0.036 |
| **Otu251** | Unclassified *Chitinophagaceae* | Environmental | | | 0.155 | 0.036 |
| **Otu120** | *Veillonella* | Human | | | 0.156 | 0.034 |
| **Otu113** | *Herbaspirillum* | Environmental | | | 0.157 | 0.033 |
| **Otu138** | *Perlucidibaca* | Environmental | | | 0.157 | 0.033 |
| **Otu146** | *Granulicatella* | Human | | | 0.161 | 0.029 |
| **Otu131** | *Actinomyces* | Both | | | 0.162 | 0.028 |
| **Otu159** | *Pseudoxanthomonas* | Environmental | | | 0.163 | 0.027 |
| **Otu094** | *Pseudomonas* | Both | | | 0.164 | 0.026 |
| **Otu075** | *Wautersiella* | Environmental | | | 0.175 | 0.017 |
| **Otu115** | *Micrococcus* | Both | | | 0.176 | 0.017 |
| **Otu072** | *Massilia* | Both | | | 0.177 | 0.016 |

| | | | | |
|---|---|---|---|---|
| **Otu022** | *Acinetobacter* | Both | 0.178 | 0.016 |
| **Otu119** | *Stigmatella* | Environmental | 0.178 | 0.015 |
| **Otu078** | *Paracoccus* | Environmental | 0.180 | 0.015 |
| **Otu091** | *Aeromicrobium* | Environmental | 0.181 | 0.014 |
| **Otu166** | *Arthrobacter* | Environmental | 0.184 | 0.012 |
| **Otu124** | *Moraxella* | Human | 0.186 | 0.012 |
| **Otu055** | *Pseudomonas* | Both | 0.189 | 0.010 |
| **Otu034** | *Janibacter* | Environmental | 0.189 | **0.010** |
| **Otu043** | *Kocuria* | Both | 0.196 | **0.008** |
| **Otu029** | *Brachybacterium* | Human | 0.202 | **0.006** |
| **Otu155** | *Tistrella* | Environmental | 0.210 | **0.004** |
| **Otu054** | *Corynebacterium* | Human | 0.213 | **0.004** |
| **Otu059** | *Luteimonas* | Environmental | 0.217 | **0.003** |
| **Otu095** | *Nocardioides* | Environmental | 0.227 | **0.002** |
| **Otu048** | *Veillonella* | Human | 0.232 | **0.001** |
| **Otu127** | *Nocardioides* | Environmental | 0.242 | **0.001** |
| **Otu046** | *Paracoccus* | Environmental | 0.244 | **0.001** |
| **Otu024** | *Acinetobacter* | Both | 0.249 | **0.001** |
